# The impacts of drift and selection on genomic evolution in holometabolous insects

**DOI:** 10.1101/072512

**Authors:** K. Jun Tong, Sebastián Duchêne, Nathan Lo, Simon Y. W. Ho

## Abstract

Genomes evolve through a medley of mutation, drift, and selection, all of which act heterogeneously across genes and lineages. The pacemaker models of genomic evolution describe the resulting patterns of evolutionary rate variation: genes that are governed by the same pacemaker exhibit the same pattern of rate heterogeneity across lineages. However, the relative importance of drift and selection in determining the structure of these pacemakers is unknown. Here, we propose a novel phylogenetic approach to explain the formation of pacemakers. We apply this method to a genomic dataset from holometabolous insects, an ancient and diverse group of organisms. We show that when drift is the dominant evolutionary process, each pacemaker tends to govern a large number of fast-evolving genes. In contrast, strong negative selection leads to many distinct pacemakers, each of which governs a few slow-evolving genes. Our results provide new insights into the interplay between drift and selection in driving genomic evolution.

## Introduction

Molecular evolution proceeds by the fixation of mutations, a process that balances stochastic drift against natural selection. The relative importance of these two forces depends on population size (Ohta 1992) and on the distribution of fitness effects of new mutations (Eyre-Walker and Keightley 2007). When mutations have neither a beneficial nor detrimental impact on fitness, their fate is determined entirely by the stochastic process of genetic drift (Kimura 1968). In contrast, purifying selection removes deleterious mutations over time. Selection is more efficient in large populations, where even small differences in selection coefficients can substantially change the relative probability of any particular mutation becoming fixed (Ohta 1992). In small populations, mutations with small fitness effects behave similarly to neutral mutations, so drift tends to be more important.

Genes are subject to varying degrees of selective constraint, leading to measurable differences in evolutionary rates (gene effects). For example, functionally important protein-coding genes tend to evolve slowly because many of the encoded amino acids are under strong selective constraint (Dickerson 1971). A simple way to detect gene effects is to examine the branch lengths of the gene trees. Genes that evolve under neutral conditions are expected to yield trees with longer branches, representing a larger total amount of genetic change. In contrast, when genetic change is retarded by purifying selection, genes are expected to yield trees with shorter branches.

The relative impacts of drift and selection also vary across species, depending on population size (Ohta 1992). For example, species with small populations are expected to evolve rapidly because of the dominance of genetic drift (Ohta 1987). In addition, differences in life-history traits, such as generation time, can produce rate heterogeneity among lineages in the tree (Bromham 2009). These lineage effects can be detected using phylogenetic methods, including relaxed-clock models (Ho and Duchêne 2014). Genes that share the same pattern of branch lengths, indicating that they are subject to the same lineage effects, are said to be governed by the same genomic pacemaker (Snir et al. 2012). On a genomic scale, there might be multiple pacemakers that each governs the evolution of a set of genes (Ho 2014; Snir 2014). The number and distribution of pacemakers throughout the genome can be detected by clustering gene trees according to their branch-length patterns (Duchêne and Ho 2015; Duchêne et al. 2016).

When genes are governed by different genomic pacemakers, there is an interaction between gene effects and lineage effects (Gillespie 1991; Muse and Gaut 1997). Consider two genes, *A* and *B*, sampled from two taxa, *x* and *y*. Both genes are responsible for important biological functions, such that their evolution is constrained. However, gene *A* is under stronger purifying selection in taxon *x* than in taxon *y*. Gene *B* is subject to the reverse conditions, with weaker purifying selection in taxon *x* and stronger selection in taxon *y*. As a consequence, the trees for these two genes display disparate branch-length patterns.

Interactions between gene effects and lineage effects are expected to be more common under conditions of selection, because the strength and direction of selection is unlikely to be uniform across species. Therefore, genes under strong selection are predicted to be governed by many genomic pacemakers and to yield trees with short branches. In contrast, genes evolving by drift are predicted to be governed by few genomic pacemakers and to yield trees with long branches. Under these conditions, most rate variation is due to lineage effects, such as those caused by differences in generation time. These lineage effects act on a genome-wide scale (Gillespie 1991), such that different genes share the same pattern of branch-length variation.

Therefore, our framework predicts that the structure of genomic pacemakers is associated with evolutionary rates across genes (Figure 1). This prediction can be tested by analysing genomic data using a phylogenetic approach, because drift and selection leave different signatures in the gene trees. Here we analyse 955 genes from 15 species of holometabolous insect. These insects undergo complete metamorphosis as part of their development. The superorder arose more than 350 million years ago (Tong et al. 2015) and is extraordinarily diverse: its members include those that are eusocial, parasites, long-distance migrators, and predators. They represent a large proportion of the global biomass and are responsible for the bulk of ecological functions on land. We statistically assigned each gene to one of a set of genomic pacemakers based on its pattern of branch-length variation. Our analyses of these data confirm the predictions of our model, whereby the number of genes governed by a pacemaker can be predicted by the evolutionary rate of those genes.

**Figure 1.**
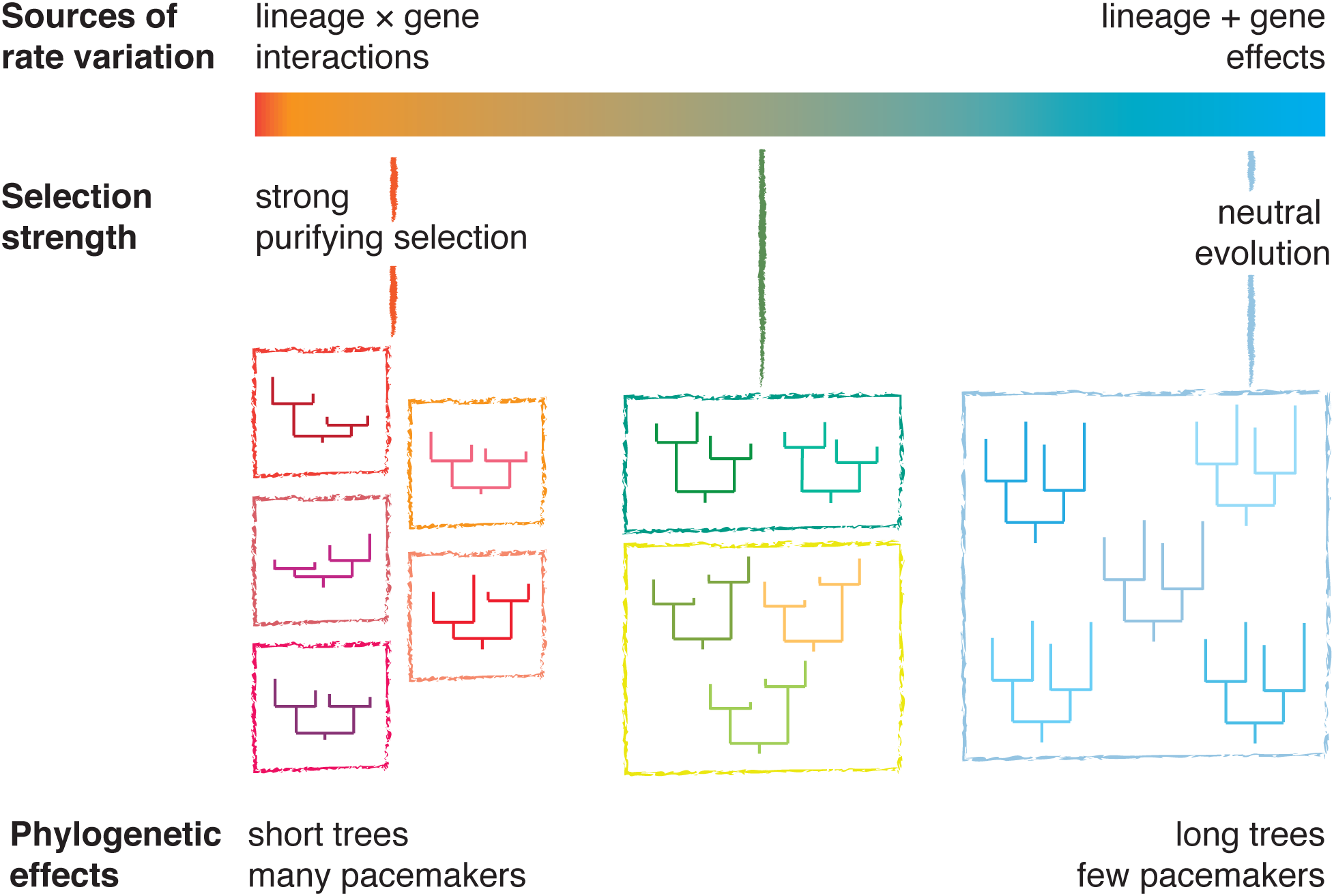
A diagram showing the relationship between evolutionary rate and pacemaker structure. Genes that are under strong purifying selection have low rates of evolution, producing short phylogenetic trees. Genes that are under weak selection are primarily subject to genetic drift and have long phylogenetic trees. Genes share patterns of among-lineage rate variation if they share similar evolutionary pressures over time and exhibit the same lineage effects. These genes are said to be governed by the same genomic pacemaker. We posit that genes whose evolution has been dominated by drift will be governed by only a small number of pacemakers, but that genes under strong purifying selection will be governed by many separate pacemakers. Genes that are under weak selection are primarily influenced by lineage effects, which act uniformly across genomes. Genes under strong selection experience gene-by-lineage interactions, which lead to distinct patterns of among-lineage rate variation across genes.

## A phylogenetic approach

We used maximum likelihood to infer the phylogeny of 15 species of holometabolous insect from a concatenated alignment of 955 genes. Based on this estimate of the tree topology, we optimized the branch lengths for each gene. Thus, the resulting gene trees shared the same topology but had their own sets of branch lengths.

We then tested the assumption that evolutionary rates are associated with the strength of negative selection. To determine the relative average rate in each gene, we took the sum of the expected number of substitutions along all of the branches in the corresponding gene tree (i.e., the tree length). Rapidly evolving genes yield longer trees, and these genes are expected to be under the weakest selective constraints. We confirmed this link by comparing gene-specific ratios of radical and conservative amino acid substitutions with the lengths of the corresponding gene trees. Our method of identifying radical and conserved substitutions is similar to that of Zhang (2000). We used a model-free, non-parametric approach to estimate this ratio. This statistic has a similar interpretation to the Kr/Kc ratio (Zhang 2000), but the absolute values are expected to be different because Kr/Kc is estimated using an explicit substitution model and phylogenetic tree. Although our method is biased towards radical substitutions, with a consequent skew in our results, it provides a fast estimate of the degree of selection (Figure 2).

**Figure 2.**
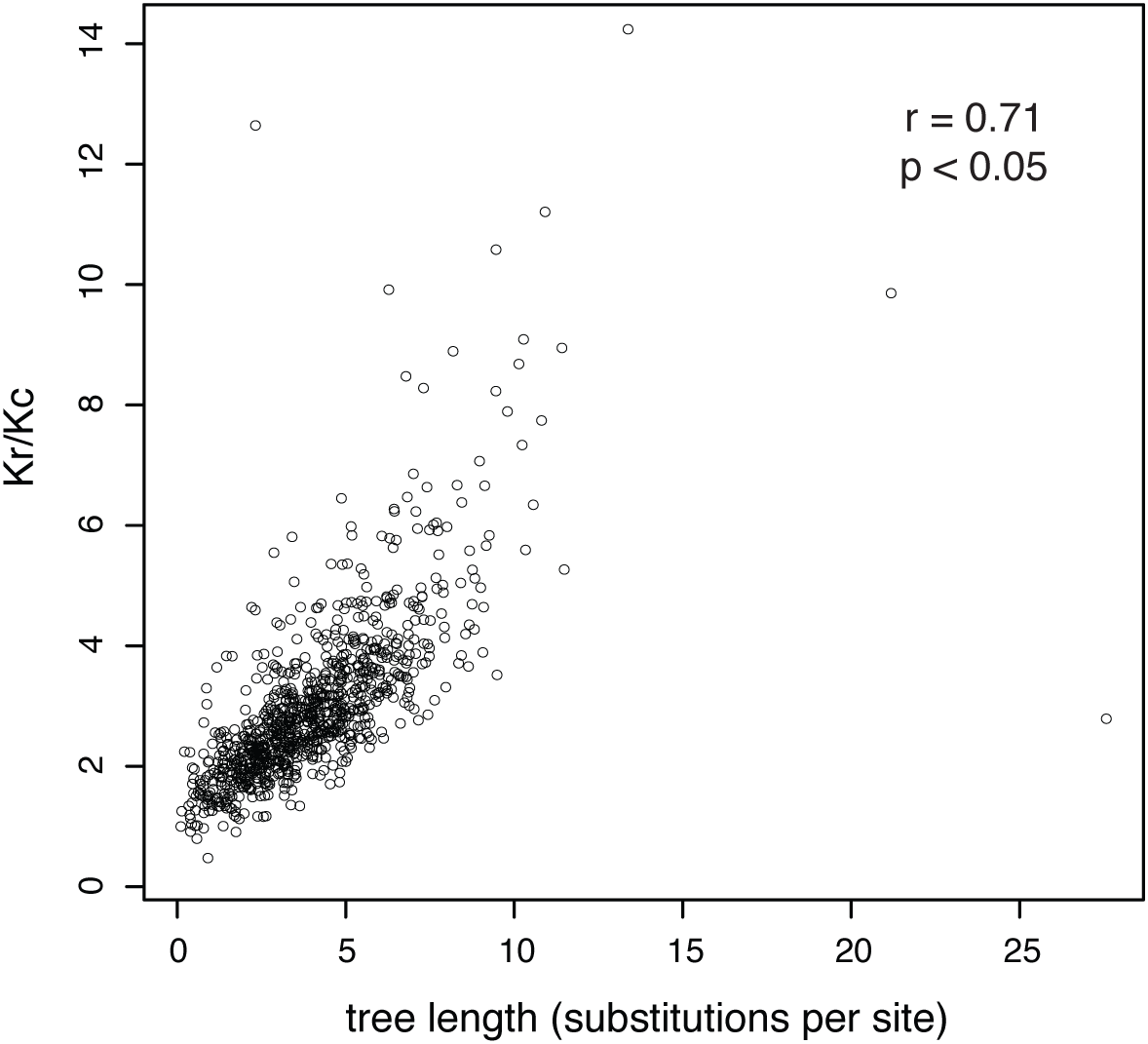
Purifying selection weakens with increasing evolutionary rate, as measured by gene-tree length. Each point represents one of 955 genes from 15 species of holometabolous insect. Kr/Kc is the ratio of estimated radical to nonradical amino acid substitutions. High Kr/Kc values indicate that radical substitutions outnumber non-radical substitutions, reflecting weak selective constraints.

Next, we wished to test the relationship between evolutionary rates and genomic pacemaker structure. To group the estimated gene trees according to their branch-length patterns, we used a clustering approach based on Gaussian mixture models, as implemented previously (Duchêne et al. 2016). Importantly, this method clusters the gene trees by their pattern of branch lengths (lineage effects), but not their overall relative evolutionary rate (gene effects). Each of the clusters that are identified using this approach represents a set of genes that are governed by the same genomic pacemaker (Duchêne and Ho 2015). In this study, we did not aim to find the optimal number of clusters for the data; instead, we wished to test our hypothesis using different numbers of clusters. Therefore, we compared the results obtained using three numbers of pacemaker clusters: 20, 30, and 40.

The gene trees were ranked according to length and divided into deciles. For each tree-length decile, we identified the number of genomic pacemakers that were represented (Figure 3). The results of our analyses confirmed our prediction of a relationship between evolutionary rate, as represented by tree length, and the structure of genomic pacemakers. Specifically, slowly evolving genes are governed by many pacemakers, whereas rapidly evolving genes are governed by few pacemakers. We found very few genes that both evolved slowly and were governed by the same genomic pacemaker. Genes that do evolve under these conditions are probably important housekeeping genes, such as those that encode histone or ribosome proteins. These genes would evolve similarly across different species, but with a very low substitution rate because they would be under strong purifying selection.

**Figure 3.**
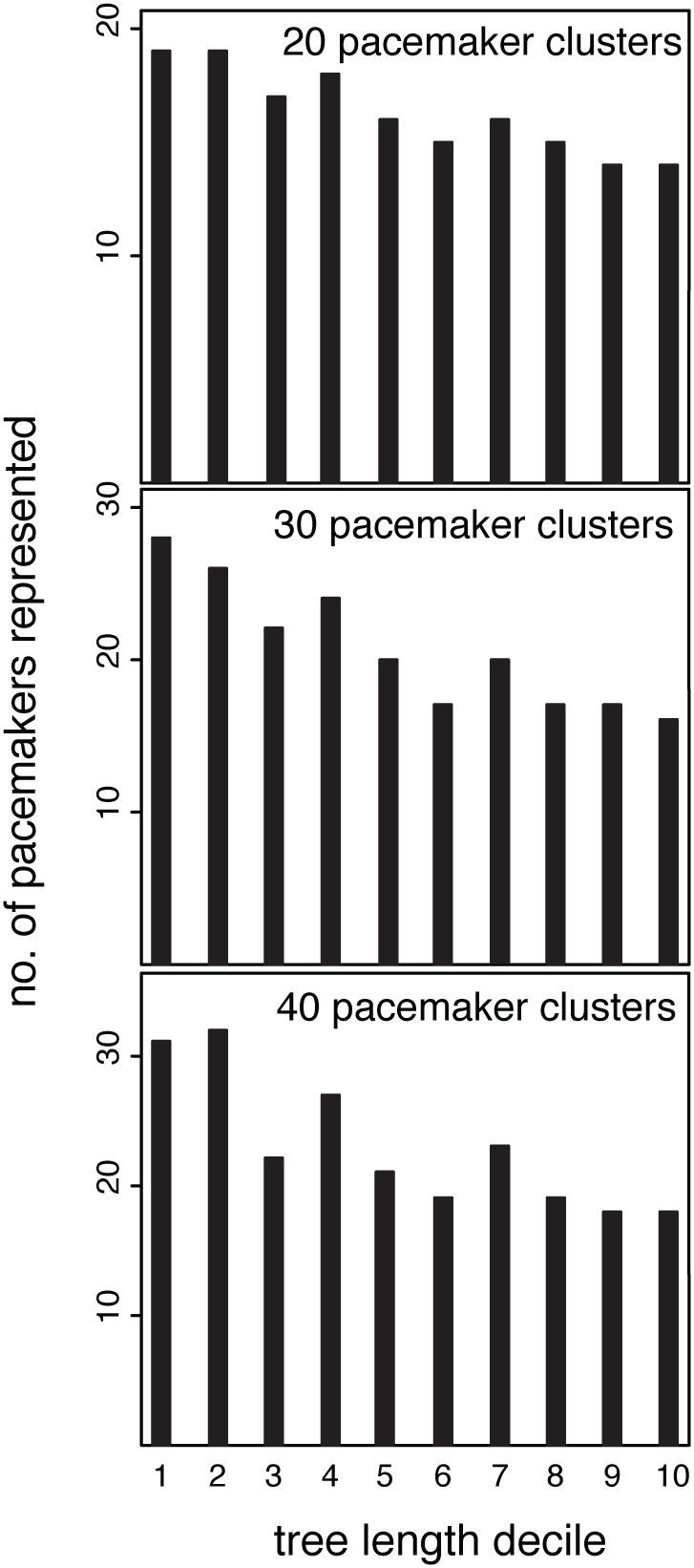
Genes with the longest trees are governed by fewer pacemakers than the decile of genes with the shortest trees. Here, genes have been sorted into deciles according to their tree lengths. Tree length is measured in substitution per site and reflects the relative rate of molecular evolution that has been experienced by a gene. For three separate pacemaker clustering schemes (20, 30, and 40), fewer pacemakers are represented in deciles of genes with higher rates.

In addition to testing the role of evolutionary rate, we investigated whether pacemaker structure could be explained by gene function. Our dataset is poorly annotated, which is typical of large datasets generated by high-throughput sequencing. This limited the scope of our investigation to enzymes because enzyme commission (EC) numbers were available for only a subset of our data. EC numbers refer to particular catalytic processes that are enabled by the enzymes. These classifications were available for 297 genes in our data set, but other genes either had incomplete annotations or did not encode enzymes. We looked at the number of pacemakers represented for each of six EC numbers. To correct for an imbalance in the number of genes within each EC category, we divided the number of represented pacemakers by the number of genes. We found that isomerase genes (EC number 2) are more likely to be governed by the same pacemaker than the genes assigned to other EC numbers (Figure 4). In contrast, transferase genes (EC number 5) are represented across many pacemakers. Interestingly, isomerases are more likely to evolve new functions in different EC classes (Martinez Cuesta et al. 2014). Such isomerase sequences might possess latent potential for selection (Dykhuizen and Hartl 1980), whereby long periods of drift produce a stream of raw genetic variation that can be subject to selection under particular conditions (Ohta 1987; 1992). We speculate that if this is the case, selection is probably occurring at the secondary or tertiary level of protein structure because the trees of the isomerase genes cluster in few pacemakers, indicating that they are subject to little selection pressure at the sequence level.

**Figure 4.**
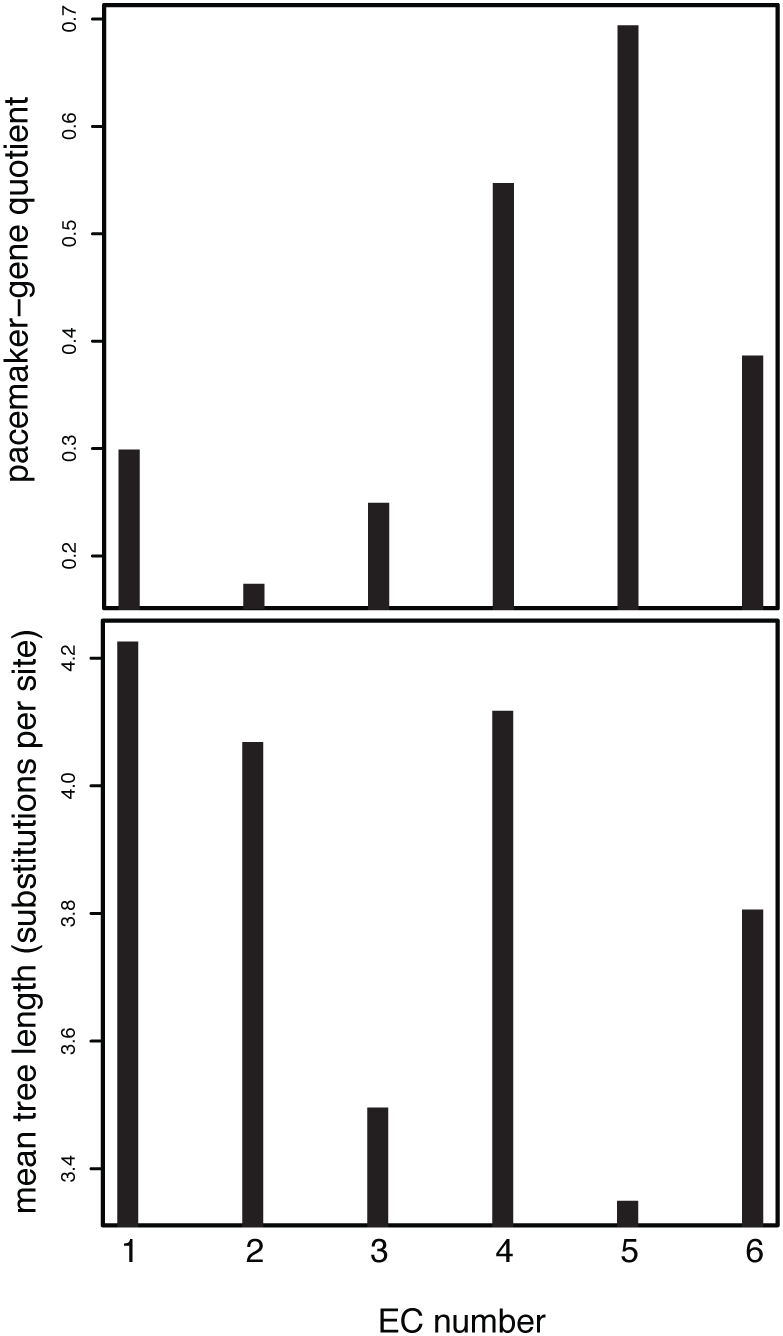
Relationship between enzyme classification (EC) number and pacemakers for 297 genes from holometabolous insects. Each EC number represents a collection of genes that share a common enzyme function. EC number 5, denoting genes that code for transferases (enzymes that move chemical functional groups from one molecule to another), is the most degenerate of categories, meaning that these genes are dispersed across more pacemakers compared to genes belonging to other EC categories. EC number 2 is the least degenerate. EC number 2 denotes isomerases, which are enzymes that convert molecules from one isomer to another. The results are based on a 40-cluster pacemaker scheme.

Our investigation of the relationship between enzyme function and pacemaker structure is limited in its statistical power. Despite our correction for the imbalance in the number of genes represented across the six EC categories, three of the six categories have 13 or fewer genes; these relatively small groups of genes might have had a large bearing on the results (Figure 4). Further clouding any signal in the dataset is the fact that some enzymes can accumulate many nucleotide changes while preserving gene function, and that gene function can change without substantial alterations to the nucleotide sequence (Martinez Cuesta et al. 2014). Enzymes can also exhibit ‘promiscuity’, whereby they evolve to catalyse new suites of reactions in addition to their normal functions (O’Brien and Herschlag 1999, Duarte et al. 2013). This hazy correspondence between nucleotide changes (or lack thereof) and biological function, the ultimate target of selection, is likely to be a contributor to the statistical noise in our pacemaker analyses. Amino acid substitutions are also thought to be more insensitive to generation-time effects compared with nucleotide substitutions, particularly nucleotide changes occurring in non-coding regions, because proteins are more likely to be targets of selection (Ohta 1992).

Finally, we fitted a random forest classifier to test whether the tree length, ratio of radical and conserved amino acid substitutions, or EC number could predict the pacemaker assignments of the genes (Liaw and Wiener 2002). Predictive power was quantified using Gini coefficients. We found that the length of the gene tree has the best predictive accuracy, with a Gini coefficient of 64. This was followed by our Kr/Kc ratio and the EC number, with respective coefficients of 60 and 25. However, the classifier has overall low predictive accuracy, suggesting that more gene features might need to be considered to provide a comprehensive mechanistic model for pacemaker assignment. This can be improved in the future with further progress in genome annotation.

## Evolutionary rates and genomic pacemakers

Our analyses reveal the roles of selection and drift in determining the structure of pacemakers across the genome. Genes that are the most weakly selected are subject to the vagaries of drift, and they tend to have the highest evolutionary rates across the genome. The main driver of rate heterogeneity in these genes is lineage effects, which explains our finding that large groups of rapidly evolving genes are governed by the same pacemakers. The most well studied lineage effect is that of differences in generation time (e.g., Thomas et al. 2010; Weller and Wu 2015). Generation time has a negative relationship with evolutionary rate because genome replication occurs more infrequently in species with long generations than in those with short generations. This is tempered by the fact that long-lived species tend to have small populations, where drift is the dominant driver of molecular evolution and leads to a higher evolutionary rate (Ohta and Kimura 1971). However, theoretical examinations suggest that certain mutagenic conditions allow the fixation of neutral mutations to be independent of population size (Welch et al. 2008).

A key problem in our attempt to describe the structure of genomic pacemakers here is the effect of fluctuating selection pressures over time. The fitness effects of mutations can vary through time, with the potential for selection that is realized under new environmental and ecological conditions (Dykhuizen and Hartl 1980; Ohta 1992). The converse might also be true: as selection dynamics shift, the magnitude of selection acting upon a gene might vary over time, or become effectively zero. Nevertheless, our phylogenetic approach is able to detect an underlying signal through this noise. The pacemaker structure that we observe here reflects groups of genes that share the same temporal patterns of rate variation. The pacemakers represented in the most rapidly evolving decile of genes might differ from one another by the shifting balance between selection and drift that has occurred over time. For instance, two pacemakers might have experienced the same total amount of evolutionary change due to drift, but differ in the periods of time in which they were subject to selection and drift, thereby generating different branch-length patterns. Our results suggest that in the pacemakers that govern the most rapidly evolving genes, these sources of fluctuation are genome-wide factors. Among the most slowly evolving genes, there is a variety of governing pacemakers because of gene-by-lineage interactions that lead to highly heterogeneous evolutionary rates.

## Implications for phylogenomic analysis

Identifying the role of evolutionary rates in the structuring of genomic pacemakers provides some useful insights into how genome-scale data might be handled in phylogenetic analysis. There is a need for new analytical methods to extract phylogenetic and temporal signals from genome-scale data without creating excessive computational demands (Ho 2014; Kumar and Hedges 2016). One promising new approach involves data-clustering to identify subsets of genes that share similar evolutionary characteristics (Duchêne et al. 2014; Mirarab et al. 2014). These techniques have already been used in phylogenomic analyses of mammals (dos Reis et al. 2012), birds (Jarvis et al. 2014), and insects (Misof et al. 2014).

Our results suggest that any form of clustering that groups genes according to their branch-length patterns will identify large groups of rapidly evolving genes and many small groups of slowly evolving genes. The latter are especially useful for studying ancient divergences because they have experienced less saturation, but they also display greater variation across genes in terms of their among-lineage rate heterogeneity. Therefore, understanding pacemaker structure has important practical implications for evolutionary dating using molecular clocks. Any molecular dating study must be based on a compromise between selecting genes with an appropriate rate of evolution, and selecting genes to minimize the variation in patterns of among-lineage rate heterogeneity.

In summary, our analysis of holometabolous insects has revealed an underlying structure to the complexity of genome evolution. Specifically, we have identified support for a model of genomic evolution in which drift and selection lead to predictable patterns of rate variation. Further detailed annotation of genomes will open the way for deeper insights into the impacts of gene function on shaping phylogenetic information. We hope that our results will spur the discovery of other widespread patterns in genome evolution and lead to improvements in phylogenomic analysis.

## Materials and Methods

We analysed a data set comprising the amino acid sequences of 955 genes from 15 insect taxa (Supplementary Table S1): two bees (*Apis mellifera* and *Bombus terrestris*), two ants (*Linepithema humile* and *Pogonomyrmex barbatus*), a wasp (*Nasonia vitripennis*), three mosquitoes (*Anopheles gambiae*, *Aedes aegypti*, and *Culex quinquefasciatus*), three flies (*Drosophila melanogaster*, *Drosophila persimilis*, and *Drosophila sechellia*), a beetle (*Tribolium castaneum*), the silkworm (*Bombyx mori*), a louse (*Pediculus humanus*), and an aphid (*Acyrthosiphon pisum*). This was modified from the data collected by Peters et al. (2014). The data were filtered to produce a subset of 955 sequences without missing data.

The tree for each gene was inferred using maximum likelihood in RAxML v8.1 (Stamatakis 2014). The same substitution model, GTR+G with four categories of site rates, was used for all genes. We ran ten replicates of each search and chose the tree with the highest likelihood score. Because we were interested in the relationship between tree length and branch-length patterns, our analyses required the topologies of the gene trees to be mutually congruent. We checked for any substantial differences in topologies between gene trees by clustering them using the k-means Partitioning Around Medoids (PAM) algorithm. We found strong support for a single cluster of tree topologies, whereby every gene supported the same set of evolutionary relationships among the 15 insect species. Accordingly, we inferred the maximum-likelihood tree from a data set comprising the 955 genes in concatenation. This tree topology was then fixed for subsequent optimization of the branch lengths for each gene tree in RAxML.

We applied a Gaussian mixture model (GMM) clustering algorithm from the Python machine learning toolkit, Scikit-learn (Pedregosa et al. 2011), which allows us to select the number of clusters into which we fit our phylogenetic data. GMM algorithms assign data to multivariate normal components and appear to work well when used to identify genomic pacemakers (Duchene et al. 2016). Based on the gene trees inferred using RAxML, we sorted the trees into deciles according to their lengths. We identified the number of different pacemakers represented in each decile, and plotted these in a histogram.

To evaluate the degree of selection acting on each gene, we calculated the ratio of radical to conserved amino acid substitutions using a custom program written in R v3.2.3 (R Core Team 2015). These values were then plotted against the corresponding gene-tree lengths. We have uploaded this program to GitHub (github.com/sebastianduchene).

We isolated a subset of 297 enzyme-coding genes from our 955-gene dataset. We matched an EC number from the NCBI database of *Drosophila melanogaster* to each of these enzyme sequences. For each EC number, we plotted the number of unique pacemakers and the mean tree length.

We investigated whether a set of variables could predict cluster assignment for the different genes. To do this, we used a random forest classifier, where the cluster assignment was the target variable and the predictors were tree length, Kr/Kc, and EC number. This method is appropriate because it does not make parametric assumptions and it is more robust to over-fitting than regression methods (Hastie et al. 2001).

## Acknowledgements

This work was supported by the Australian Research Council (grant number DP160104173). KJT was supported by an Australian Postgraduate Award.

